# A Bayesian model of acquisition and clearance of bacterial colonization incorporating within-host variation

**DOI:** 10.1101/429464

**Authors:** Marko Järvenpää, Mohamad R. Abdul Sater, Georgia K. Lagoudas, Paul C. Blainey, Loren G. Miller, James A. McKinnell, Susan S. Huang, Yonatan H. Grad, Pekka Marttinen

## Abstract

Bacterial populations that colonize a host can play important roles in host health, including serving as a reservoir that transmits to other hosts and from which invasive strains emerge, thus emphasizing the importance of understanding rates of acquisition and clearance of colonizing populations. Studies of colonization dynamics have been based on assessment of whether serial samples represent a single population or distinct colonization events. With the use of whole genome sequencing to determine genetic distance between isolates, a common solution to estimate acquisition and clearance rates has been to assume a fixed genetic distance threshold below which isolates are considered to represent the same strain. However, this approach is often inadequate to account for the diversity of the underlying within-host evolving population, the time intervals between consecutive measurements, and the uncertainty in the estimated acquisition and clearance rates. Here, we present a fully Bayesian model that provides probabilities of whether two strains should be considered the same, allowing us to determine bacterial clearance and acquisition from genomes sampled over time. Our method explicitly models the within-host variation using population genetic simulation, and the inference is done using a combination of Approximate Bayesian Computation (ABC) and Markov Chain Monte Carlo (MCMC). We validate the method with multiple carefully conducted simulations and demonstrate its use in practice by analyzing a collection of methicillin resistant Staphylococcus aureus (MRSA) isolates from a large recently completed longitudinal clinical study. An R-code implementation of the method is freely available at: https://github.com/mjarvenpaa/bacterial-colonization-model.git.

**Author summary:** As colonizing bacterial populations are the source for much transmission and a reservoir for infection, they are a major focus of interest clinically and epidemiologically. Understanding the dynamics of colonization depends on being able to confidently identify acquisition and clearance events given intermittent sampling of hosts. To do so, we need a model of within-host bacterial population evolution from acquisition through the time of sampling that enables estimation of whether two samples are derived from the same population. Past efforts have frequently relied on empirical genetic distance thresholds that forgo an underlying model or employ a simple molecular clock model. Here, we present an inferential method that accounts for the timing of sample collection and population diversification, to provide a probabilistic estimate for whether two isolates represent the same colonizing strain. This method has implications for understanding the dynamics of acquisition and clearance of colonizing bacteria, and the impact on these rates by factors such as sensitivity of the sampling method, pathogen genotype, competition with other carriage bacteria, host immune response, and antibiotic exposure.

## Introduction

Colonizing bacterial populations are often the source of infecting strains and transmission to new hosts [1–5], making it important to understand the dynamics of these populations and the factors that contribute to persistent colonization and to the success or failure of clinical decolonization protocols. The study of colonization dynamics is based on inferring whether bacteria from samples collected over time represent the same population or distinct colonization events, thereby permitting calculation of rates of acquisition and clearance [6, 7]. Whole genome sequencing has provided a detailed measure of genetic distance between isolates, which can then be used to infer the relationship between them [8–11]. While to date most studies have used genetic distance thresholds as the basis for determining the relationship between isolates [8, 10], here we improve on these heuristic strategies and present a robust and accurate fully Bayesian model that provides probabilities of whether two strains should be considered the same, allowing us to determine bacterial clearance and acquisition from genomes sampled over time.

An example of a typical individual-level longitudinally sampled data set from a study population is shown in Fig 1: each ’row’ represents a patient, x-axis is time, and dots are the genomes sampled at multiple time points. Dot color refers to different, easily distinguishable, sequence types (ST). The coloured number between two consecutive samples reflects the distance between the genomes, and we see that even within the same ST the distances may vary considerably, and, therefore, determining whether the changes can be explained by within-host evolution only, is challenging. Intuitively, if two genomes are very similar, we interpret this as a single strain colonizing the host. On the other hand, two very different genomes, even if the same ST, are interpreted as two different strains, obtained either jointly or separately as two acquisitions. With these data, we would like to address questions including: to what extent are people persistently colonized, cleared, and recolonized? If recolonized, what is the likelihood that it is the same or a distinct strain? To address these questions, previous works have relied on using a threshold number of single nucleotide polymorphisms (SNPs) to define a strain. Optimally, however, the SNP distance between the genomes observed and the interval between the sampling time defines a probability that the two genomes represent the same strain. Such data are critical for understanding within-host dynamics, response to interventions, and transmission.

**Fig 1.**
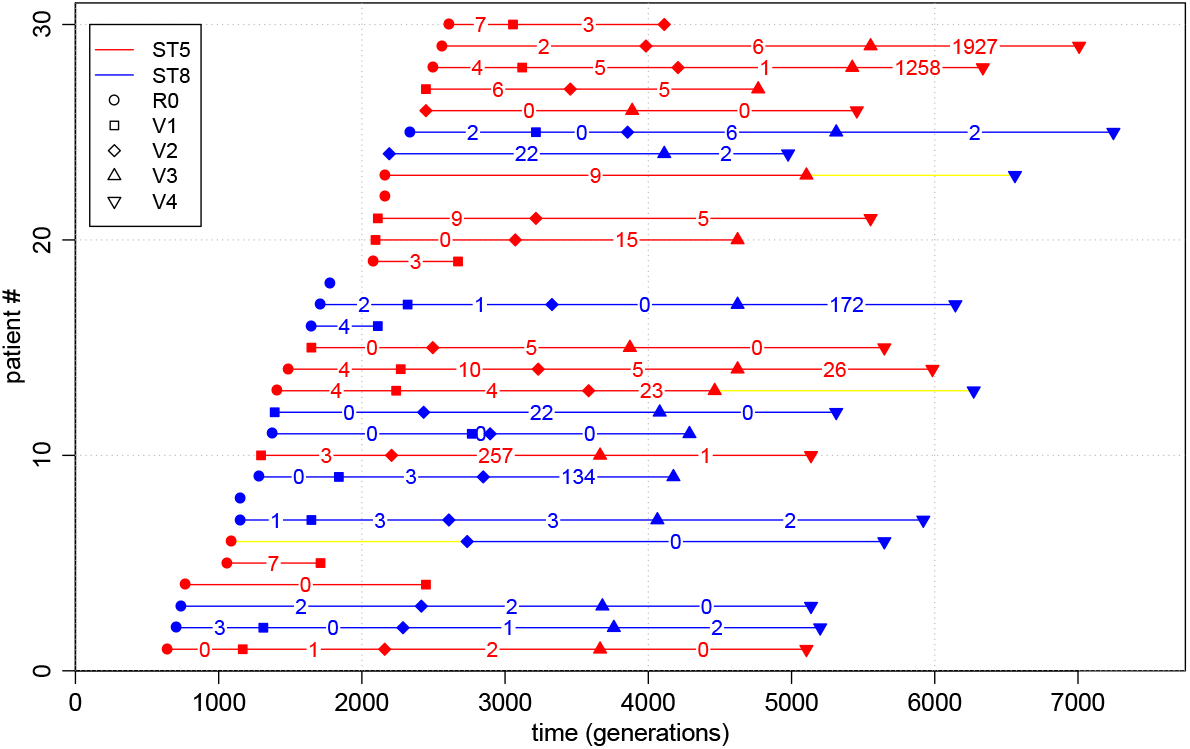
Illustration of a subset of the data used in the study. Each row corresponds to one patient and only the first 30 patients are shown. R0 is the initial hospital visit and V1, V2 etc. are the further visits. Red colour refers to ST5 and blue to ST8 and the coloured numbers are the amount of mutations *d_i_*. Yellow colour highlights the cases where the ST changes from ST 5 to ST 8.

Previously, transitions between different colonizing bacteria have been modeled using hidden Markov models [12] with states corresponding to different colonizing STs. However, this approach is not suitable for modeling within a single ST, where acquisition and clearance must be determined based on a small number of mutations. Crucial for interpreting such small differences is a model for within-host variation [8, 13], specifying the number of mutations expected by evolution within the host. Population genetic models can be used for understanding the variation in an evolving population [14]. A major difficulty in fitting such models to data like those shown in Fig 1 is that the information contained by the data is extremely limited regarding the variation within the host: a single time point is summarized with just a single (or a few) genomes, and must serve to represent the whole within-host population. While some studies use genome sequence from multiple isolates to achieve a more complete characterization of within-host diversity [3, 10], these tend to be limited in terms of the number of time points and/or patients.

The Bayesian statistical framework can be used to combine information from multiple data sources. In the Bayesian approach, a prior distribution is updated using the laws of probability into a posterior distribution in the light of the observations, and this can be repeated multiple times with different data sets [15, 16]. Approximate Bayesian computation (ABC) is particularly useful with population genetic models, where the likelihood function may be difficult to specify explicitly, but simulating the model is straightforward [17, 18]. ABC has recently been introduced in bacterial population genetics [19–22]. Here, we present a Bayesian model for determining whether two genomes should be considered the same strain, enabling a strategy grounded in population genetics to make inferences about acquisition and clearance from data of closely related genomes. Benefits of the fully Bayesian analysis include: rigorous quantification of uncertainty, explicit statement of modeling assumptions (open for criticism and further development when needed), and straightforward utilization of multiple data sources. We demonstrate these benefits by analyzing a large collection longitudinally collected methicillin resistant *Staphylococcus aureus* (MRSA) genomes, obtained through a clinical trial (Project CLEAR) to evaluate the effectiveness of an MRSA decolonization protocol [23]. This method for identifying strains with explicit assessment of uncertainty will enable studies of the characteristics–both host and pathogen–that impact colonization in the presence and absence of interventions.

## Methods

### Overview of the model

One input data item for our model consists of a pair of genomes that are of the same ST, sampled from the same individual at two consecutive time points (or possibly with an intervening time point with no samples or a sample of a different ST). Each of these data items (i.e. pairs of consecutive genomes) is summarized in terms of two quantities: the distance between the genomes and the difference between their sampling times (see Fig 1). Hence, the observed data *D* can be written as consisting of pairs (*d_i_, t_i_*), *i* = 1,…, *N*, where *t_i_* > 0 is the time between the sampling of the genomes, *d_i_* ∈ {0, 1, 2, …} is the observed distance, and *N* the total number of genome pairs that satisfy the criteria (i.e. same patient, same ST, consecutive time points or possibly with an intervening time point with no samples or a sample of a different ST). The restriction to genome pairs of the same ST stems from the fact that different STs will always be considered different strains anyway.

There are two possible explanations for the observed distances. If the genomes are from the same strain, we expect their distance to be relatively small. If the genomes are from different strains, we expect a greater distance. Below we define two probabilistic models that represent these two alternative explanations. These models are then combined into one overall mixture model, which assumes that the distance between a certain pair of genomes is generated either from the ’same strain’ model or the ’different strain’ model, and enables calculation of the probabilities of these two alternatives for each genome pair, rather than relying on a fixed threshold to distinguish between them.

An essential part of our approach is a population genetic simulation which allows us to model the within-host variation, and hence make probabilistic statements of the plausibilities of the ’same strain’ vs. ’different strain’ models. For this purpose, we adopt the common Wright-Fisher (W-F) simulation model, see e.g. [24], with a constant mutation rate and population size, which are estimated from the data. The simulation is started with all genomes being the same, which corresponds to a biological scenario according to which a colonization begins with a single isolate multiplying rapidly until reaching the maximum ’capacity’, followed by slow diversification of the population. This assumption is supported by the fact that in the distance distribution, in cases where the acquisition time was known and had happened recently, very little variation was observed in the population. See the Discussion section for more details on the modeling assumptions. Overview of the approach, including data sets, models, and methods for inference, is outlined in Fig 2 and discussed below in detail.

**Fig 2.**
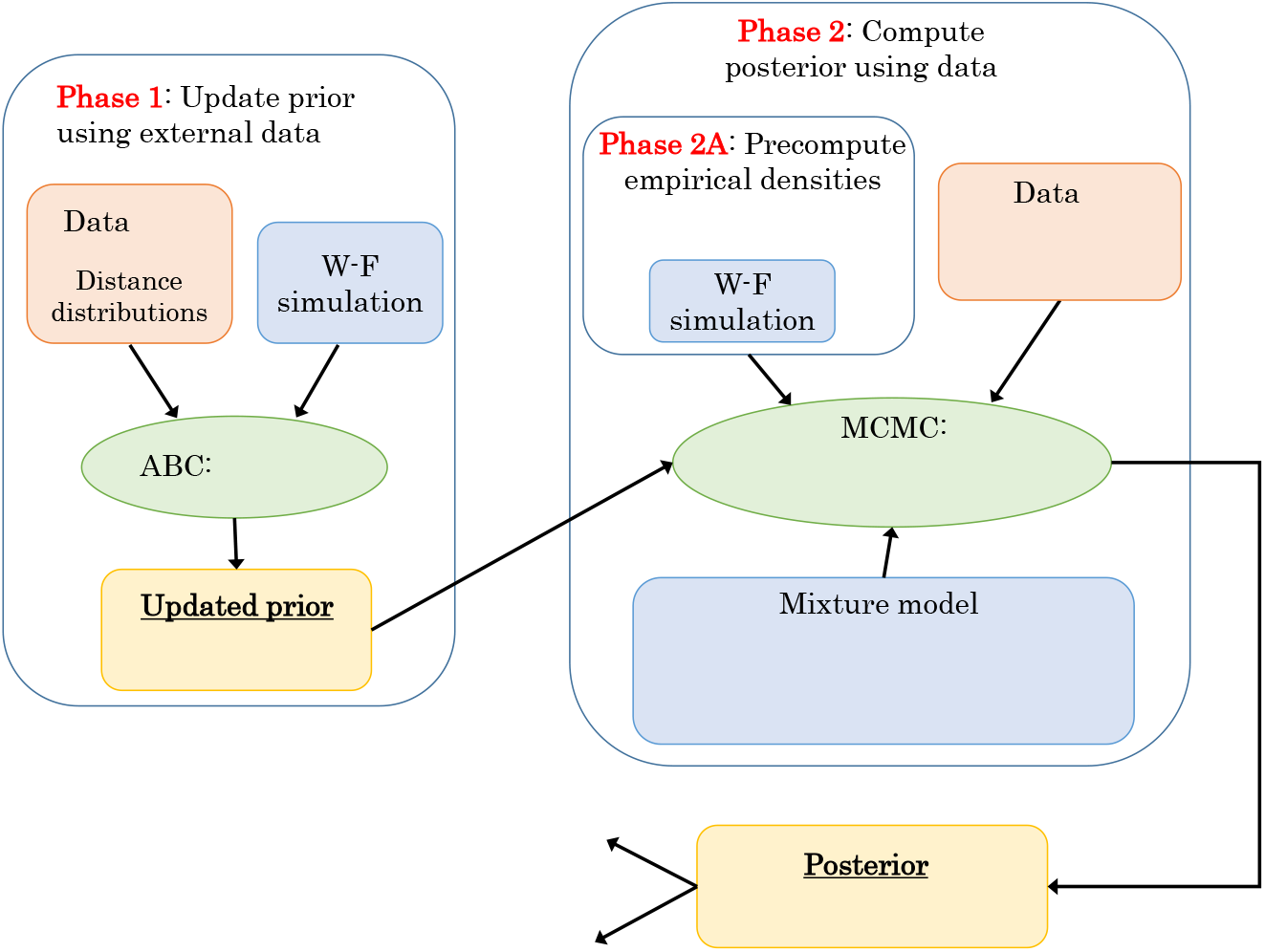
Overview of the modeling and data fitting steps. In Phase 1 we update our prior information on parameters (*n*_eff_; *μ*) based on external data *D_0_*. In phase 2 we estimate all the parameters of the (mixture) model using MCMC, precomputed distance distributions *pS* and the information obtained in Phase 1. The fitted model can be used to e.g. obtain the same strain probability for a new (future) measurement.

### Model *pS*: Same strain

Let (*s_i_*_1_, *s_i_*_2_) denote a pair of genomes with distance *d_i_*, sampled from a patient at two consecutive time points (see the previous section) with time *t_i_* between taking the samples. Here we present a model, i.e., a probability distribution *p_S_*(*d_i_* | *t_i_, n*_eff_, *μ*), which tells what kind of distances we should expect if the genomes are from the same strain. The parameter *n*_eff_ is the effective population size and *μ* is the mutation rate. We model *d_i_* as 
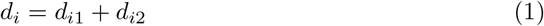
where we have defined 
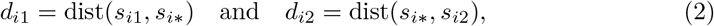
where dist(·,·) is a distance function that tells the number of mutations between its arguments, and *s_i_*_*_ is the unique ancestor of *s_i_*_2_ that was present in the host when *s_i_*_1_ was sampled, and which has descended within the host from the same genome as *s_i_*_1_(see Fig 3A). The Equation 1 is valid when mutations between *s_i_*_1_, and *s_i_*_*_, and *s_i_*_*_ and *s_i_*_2_ have occurred in different sites, which is true with a high probability when the genomes are long (millions of bps) compared to the number of mutations (dozens or a few hundred at most). The probability distribution of *d_i_*_1_ which we will denote by *p_sim_*(*d_i_*_1_|*n_eff_, μ*), and which is not available analytically and does not depend on *t_i_*, represents the within-host variation at a single time point, and we approximate it as 
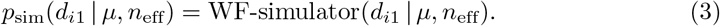

The distribution of *d_i_*_2_ is assumed to be 
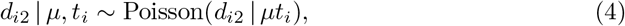
that is, mutations are assumed to occur according to a Poisson process with the rate parameter *μ*.

### Model *p_D_*: Different strains

Model *p_D_* represents the case that the genomes si1 and si2 are from different strains, which we define to mean that their most recent common ancestor (MRCA), denoted by *s_iA_*, resided outside the host. The time between *s_iA_* and *s_i_*_1_ is denoted by *t*_0*i*_ (see Fig 3B). Under model *p_D_*, we assume that the distribution of the distance *d_i_* is

**Fig 3.**
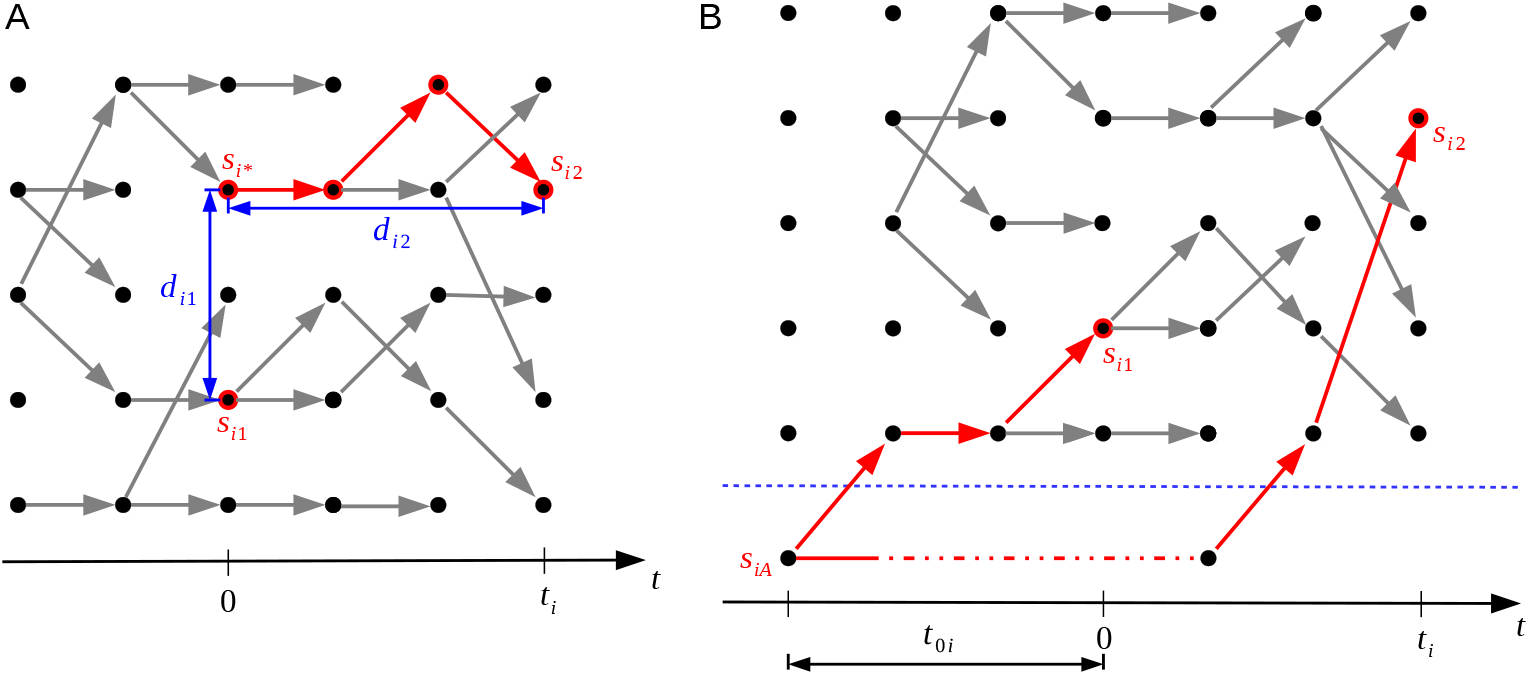
Outline of the ’same strain’ and ’different strain’ models. Model *p_D_* on the left (panel A) represents the situation where the genomes denoted by *s_i_*_1_ and *s_i_*_2_ are of the same strain. Model *p_S_* on the right (panel B) shows the case where these genomes are of different strains. Time flows from left to right in the figures, the dots represent individual genomes, and the edges parent-offspring relationships.

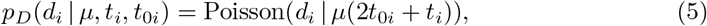
where the values of *t*_0*i*_ are unknown and will be estimated, but let us assume for now that they are known. One difference between the same strain model *p_S_* (defined by Equations 1, 3, 4) and the different strain model *p_D_* (Equation 5) is that the former uses Wright-Fisher simulation, whereas the latter does not. The reason is that the within-host variation is bounded, occasionally increasing and decreasing, which is reflected by the constant population size of the Wright-Fisher simulation in the same strain model. On the other hand, in the different strain model the distance between *s_i_*_1_ and *s_i_*_2_ can in principle increase without bound, given enough time since their common ancestor, because they diverged and evolved outside the host.

### Mixture model

With the two alternative models for the distance, we can write the full model, which assumes that each distance observation is distributed according to 
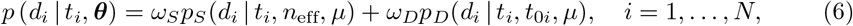
where *θ* denotes jointly all the parameters of the models, i.e., *θ* = (*n*_eff_, *μ, ω_S_, ω_D_, t*_01_,…, *t*_0*N*_). The parameter *ω_S_* represents the proportion of pairs from the same strain and *ω_D_* is the proportion of pairs from different strains, such that *ω_S_* + *ω_D_* = 1. To learn the unknown parameters θ, we need to fit the model to data, but before going into details, we discuss how to use an external data set to update the prior distribution about the mutation rate *μ* and the effective sample size *n*_eff_. This updated distribution will itself be used as the prior in the mixture model.

### ABC inference to update the prior using external data

Simulations with the W-F model are used in our approach for two purposes: 1) to incorporate information from an external data set to update the prior on the mutation rate *μ* and the effective sample size *n*_eff_, and 2) to define empirically the distribution *p_S_*(*d_i_*|*t_i_, n*_eff_, *μ*) required in the mixture model. Here we discuss the first task.

As external data we use measurements from eight patients colonised with MSSA [3], comprising nasal swabs from two time points for each patient, such that the acquisition is known to have happened approximately just before the first swab. Multiple genomes were sequenced from each sample, and the distributions of pairwise distances between the genomes provide snapshots to the within-host variability at the two time points for each individual, and these distance distributions are used as data. We exclude one patient (number 1219) because according to [3] this patient was likely infected already long before the first sample. The data set also contains observations from an additional 13 patients from [13], denoted by letters from A to M in [3]. For these patients, distance distributions from only one time point are available, and the acquisition times are unknown. The data comprising the distance distributions from the 7 patients (two time points) and the additional 13 patients (a single time point) are jointly denoted by *D*_0_.

To learn about the unknown parameters *n*_eff_ and *μ*, we first note that their values affect the distance distribution of a population resulting from a W-F simulation with the specified values (Fig. 4). To estimate these parameters, we try to find such values for them which make the output of the W-F similar to the observed distance distributions *D*_0_. Since the corresponding likelihood function is unavailable, standard statistical techniques for model fitting do not apply. Therefore, we use Approximate Bayesian Computation (ABC), a class of methods for Bayesian inference when the likelihood is either unavailable or too expensive to evaluate but simulating the model is feasible, see [17, 18, 25, 26] for an overview on ABC. The basic ABC rejection sampler algorithm for the model fitting consists of the following steps:

**Fig 4.**
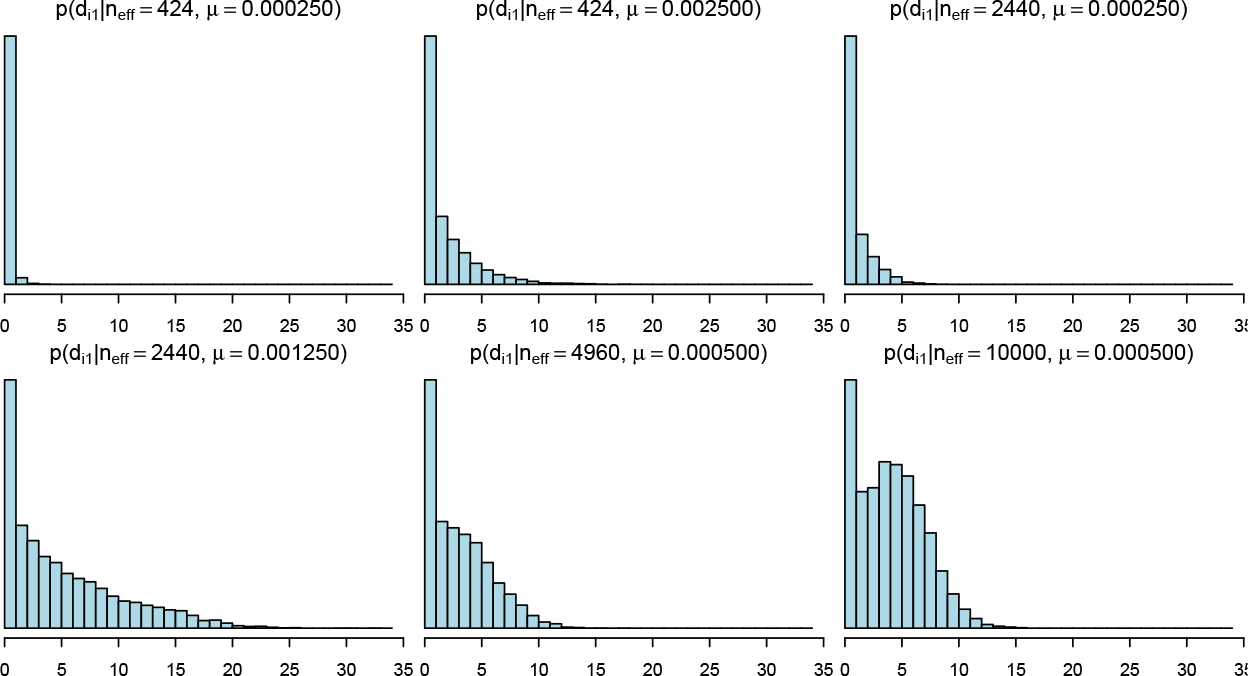
Distributions of pairwise distances for populations simulated with different parameters. The histograms show the estimated probability mass functions 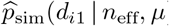 with selected parameter vectors (*n*_eff_, *μ*). Increasing *μ* and/or *n*_eff_ tends to increase the distances. The distance distribution can also be bimodal as the subfigure in lower right corner shows. Each histogram represent variability in a simulated population at a single time point 6,000 generations after the beginning of the simulation.

1. Simulate a parameter vector (*n*_eff_, *μ*) from the prior distribution *p*(*n*_eff_, *μ*).
2. Generate a pseudo-data similar to the observed data *D*_0_ by running the W-F model separately for each patient using the parameter (*n*_eff_, *μ*).
3. Accept the parameter (*n*_eff_, *μ*) as a sample from the (approximate) posterior distribution if the discrepancy between the observed and simulated data is smaller than a specified threshold *∊*.

The quality of the resulting ABC approximation depends on the selection of the discrepancy function, the threshold *∊* and the number of accepted samples. Broadly speaking, if the discrepancy summarizes the information in the data completely (e.g. it is a function of the sufficient statistics) and *∊* is arbitrarily small, the approximation error becomes negligible and the samples are generated from the exact posterior. In practice, choosing *∊* very small makes the algorithm inefficient since many simulations are needed to obtain an accepted sample even with the optimal value of the parameter. Also, finding a good discrepancy function may be difficult because sufficient statistics are typically unavailable. Many sophisticated ABC variants exist, see e.g. [18, 26] and the references therein, but as we need to estimate only two parameters (one of which is discrete) and because running the simulations in parallel is straightforward with the basic algorithm, we use a the ABC rejection sampler outlined above, with details discussed below.

In [13], MRSA evolution was simulated using parameters derived from the following estimates: 8 mutations per genome per year and generation length of 90 minutes (the whole year is thus 5840 generations). This gives mutation rate of 0.0019 per genome per generation, approximately 6.3 × 10^−10^ mutations per site per generation assuming the genome length of 3 Mbp. We also use the generation time of 90 minutes, originally derived by [13] from the estimated doubling time of *Staphylococcus aureus* [27]. We use independent uniform priors for the parameters of the W-F model, so that 
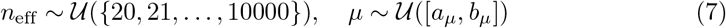
with *a_μ_* = 0.00005 and *b_μ_* = 0.005 mutations per genome per generation.

We argue that reasonable parameters should produce populations with similar histograms of the pairwise distances compared to the observations at the corresponding times. Consequently, we use the discrepancy Δ defined as 
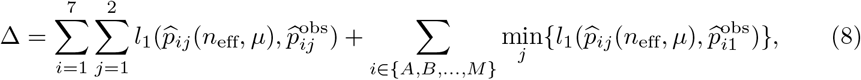
where 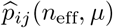 and 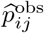 are the simulated and observed empirical distributions of pairwise distances for patient *i* with time point *j*, respectively, and *l*_1_(·,·) denotes the *L*^1^ distance between the distributions. In principle, the unknown acquisition times for the 13 patients (A-M) could be estimated by making each of them an additional parameter. However, ABC in the resulting 15 dimensional parameter space would be difficult due to the curse of dimensionality. Instead, as shown in the Eq 8, we use these data such that we supplement the unknown times with values that produce the minimum discrepancy. This way, parameters that never produce enough variability to match the observations will increase the discrepancy, allowing us to gain evidence against such unreasonable values, even if the exact times are unknown and too computationally costly to infer.

Instead of simulating (*n*_eff_, *μ*) samples from the prior we perform equivalent grid-based computations. That is, we consider an equidistant 50 *×* 50 grid of (*n*_eff_, *μ*) values and simulate the model 1, 000 times at each grid point. However, in preliminary experiments we noticed that if *n*_eff_ and *μ* are simultaneously large, the amount of mutations produced by the model increases rapidly and it is clear that the simulated pairwise distances are always greater than in the observed data, and also the computation time and memory usage become prohibitive. Thus, we do not run the full set of 1000 simulations in this parameter region because it is clear that the posterior density would be negligible. Finally, the threshold *∊* is chosen such that 5, 000 out of the total of almost 1 million simulations are below the threshold, corresponding to the acceptance probability of 0.0057.

## Details of the mixture model

We now discuss the mixture model in detail and then derive an efficient algorithm to estimate its parameters. Because the values of *t*_0*i*_ in Eq 5, denoting the times to the MRCAs in case the sequences are different strains, are unknown, we model them as random variables and give each of them a prior distribution 
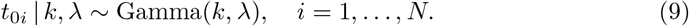
We further specify a weakly informative prior for λ such that 
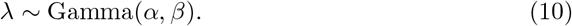

The parameter is thus shared between different *t*_0*i*_ which allows us to learn about its distribution.

If *k* = 1, then the Gamma distribution in Eq 9 reduces to the Exponential, which, however, does not reflect our prior understanding of reasonable value of *t*_0*i*_ because the mode of the resulting distribution is at zero, corresponding to a very recent common ancestor for genomes considered to be from different strains. Instead, we set *k* = 5, *α* = 2.5, and *β* = 1600, which approximately correspond to the mean and standard deviation of 5800 and 8400 generations, respectively. This weakly informative prior reflects the notion that different strains diverged on average approximately a year ago, but with a large variance. Furthermore, if the time between samples, *t_i_*, is three months, the prior translates to an expectation that, if the sampled genomes are from different strains, they are on average 30 mutations apart, with a large standard deviation of 50 mutations. Moreover, the density has a heavy tail to account for some possibly much greater distances. The formulas used to compute these values and other useful facts about the prior are provided in the supplementary material.

An equivalent way of writing the mixture model in Eq 6, which also simplifies the computations, is to introduce hidden labels which specify the component which generated each observation *d_i_*, see [28]. We thus define latent variables 
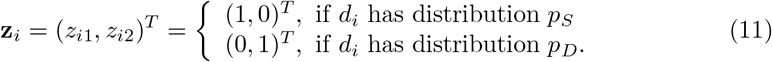

The prior density for the latent variables z is 
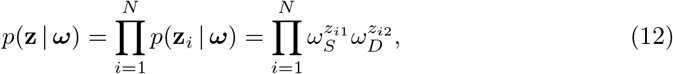
where we have used vector notation t = (*t*_1_,…, *t_N_*)^*T*^, d = (*d*_1_,…, *d_N_*)^*T*^, z = (z_1_,…, z_*N*_)^*T*^, t_0_ = (*t*_01_,…, *t*_0*N*_)^*T*^ and *ω* = (*ω_S_, ω_D_*)^*T*^. We augment the parameter θ to represent jointly all model parameters in Eq 6 and the prior densities specified in Eq 9 and 10, i.e., *θ* = (*n*_eff_, *μ, ω*, z, t_0_, λ)^*T*^. To complete the model specification, we must specify the prior for *ω, n*_eff_ and *μ*. We use 
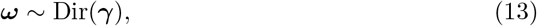
that is, a Dirichlet distribution with parameter γ = (1,1)^*T*^. We use the posterior *p*(*n*_eff_, *μ*|*D*_0_), obtained by ABC using the external data *D*_0_ as discussed in the previous section, as the (joint) prior for (*n*_eff_, *μ*).

## Bayesian inference for the mixture model

We now show how the mixture model can be fit efficiently to data. The joint probability distribution for the data d and the parameters *θ* can be now written as 
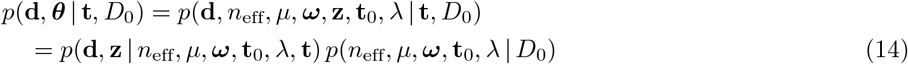

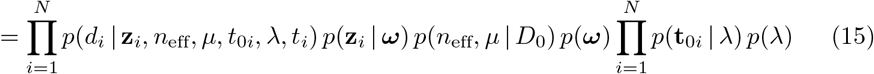
We use Gibbs sampling, which is an MCMC algorithm, to sample from the posterior density. The algorithm exploits the hierarchical structure of the model and it proceeds by iteratively sampling from the conditional density of each variable (or a block of variables) at a time [29]. In the following we derive the conditional densities for the Gibbs sampling algorithm. We observed that some of the parameters are highly correlated which causes slow mixing of the resulting Markov chain and thus inefficient exploration of the parameter space. To make the algorithm more efficient, we reparametrise the model by defining new parameters *θ*′ = (*n*_eff_, *μ, ω, z, η, λ*) via the transformation *η* = *μ*t_0_ and we use the Gibbs sampler for the transformed parameters *θ*^′^. This common strategy [29] resolves the problem arising from correlations between *t*_0*i*_ and *μ*, because the magnitudes of all *η_i_* can now be changed simultaneously by a single *μ* update. The original variables *t*_0*i*_ can be obtained from the generated samples as *t*_0*i*_ = *η_i_*/*μ*.

The joint probability in Eq 15 for the transformed parameters then becomes 
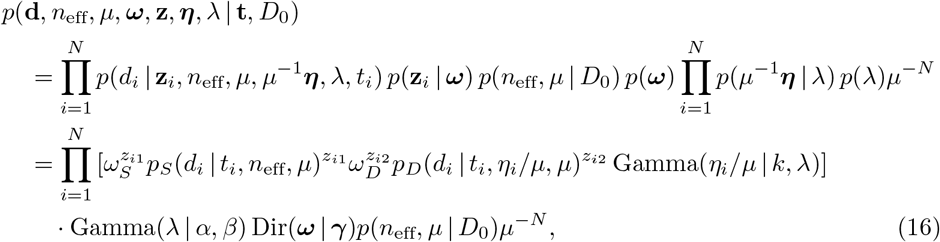
where *μ^−^N* is the determinant of the Jacobian of the inverse transformation. Computing the conditional density of parameter *ω* is straightforward. We neglect those terms in Eq 16 that do not depend on *ω* and recognise the resulting formula as an unnormalised Dirichlet distribution. We then obtain 
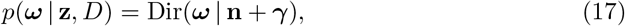
with n = (*n*_1_, *n*_2_)^*T*^, where 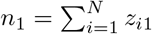 and 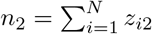. Next we consider the latent variables z_*i*_. We see that the conditional distribution of z_*i*_ for any *i* = 1,…, *N* does not depend on other latent variables z_*j*_, *j* ≠ *i*. Specifically, we obtain 
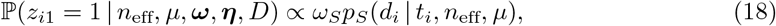

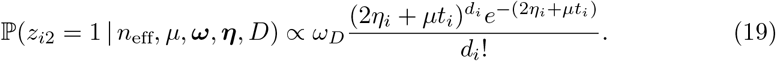
We expect the effective sample size *n*_eff_ and the mutation parameter *μ* to be correlated a posteriori so we include them to the same block and update them together. We also include λ to this block as it also tends to be correlated with *n*_eff_ and *μ*. It is convenient to replace the sampling step from *p*(*n*_eff_, *μ, λ*| *ω, z,η, D, D*_0_) with the following two consecutive sampling steps: first sample from *p*(*n*_eff_, *μ*,|*ω, z*,*η, D, D*_0_)= ∫ *p*(*n*_eff_, *μ*, λ |*ω, z*,*η, D, D*_0_) dλ and then sample from *p*(λ | *n*_eff_, *μ, ω, z*,*η, D, D*_0_). From Eq 16 we observe that 
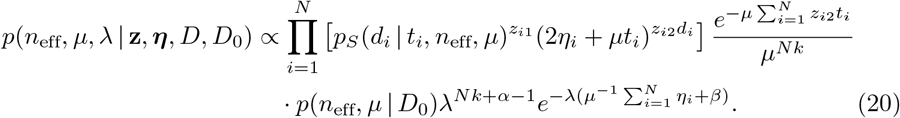

The above formula is recognised to be proportional to a Gamma density as a function of λ. We can thus marginalise λ easily to obtain the following density for the first step 
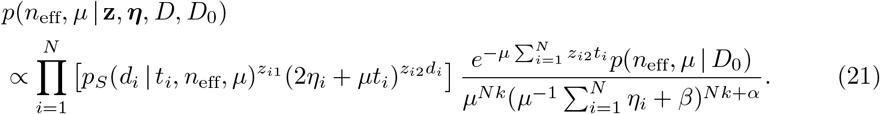

In the second step, we sample from the probability density 
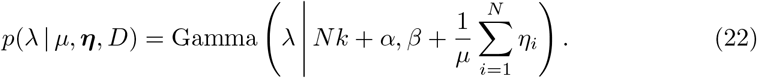

This formula follows directly from Eq 20.

Sampling from Eq 21 and sampling z using Eq 18 are challenging because *p_S_* is defined implicitly via the W-F simulation model. Consequently, we will consider an approximation that allows to compute *p_S_*(*d_i_*| *t_i_, n*_eff_, *μ*) for any proposed point (*n*_eff_, *μ*) and all values of *d_i_* and *t_i_* in the data. Since *d_i_* = *d_i_*_1_ + *d_i_*_2_, we can use the convolution formula for a sum of discrete random variables to see that 
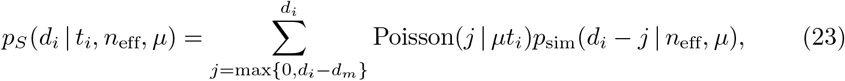
where *p*_sim_ specifies the distribution for a distance between two genomes as in Eq 3 and *d_m_* is the maximum distance that can be obtained from *p*_sim_.

Since *p*_sim_(*d_i_*_1_ | *n*_eff_, *μ*) is not available analytically, we estimate this probability mass function by simulation. A special case is if we know that there is no variation in the population at the time of taking the first sample *s_i_*_1_, which can happen if we know that the acquisition happened just before the first sample. In this case, *d_i_*_1_ = 0, and we do not need the simulation. Since this is usually not the case, we use a general solution as follows: for each (*n*_eff_, *μ*) value, we sample independently 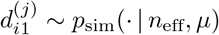 by simulating the W-F model, sample a pair of genomes at a fixed time *t* from the simulated population, and compute their distance 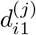. This is repeated for *j* = 1,…, *s*. Since *d_i_*_1_ is discrete, we approximate 
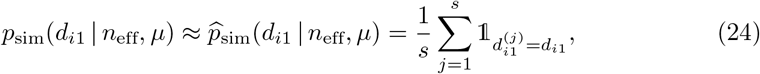
for all *i*. Since in data *D* we do not know the acquisition times, we set *t* = 6000 generations and use this same value for all *i*. This large value represents a steady state of the simulation, where the variation in the population occasionally increases and decreases as new lineages emerge and old ones die out, which can be seen as corresponding to a reasonable default expectation about population variability when the true acquisition time is unknown. While this assumption was introduced for computational necessity, it can be justified by considering its impact on the inferences: the simplification may cause slightly overestimated distances *d_i_*_1_ if many acquisitions in reality happened very recently. The consequence is that the criterion for reporting new acquisitions becomes more *conservative*, because now the ’same strain’ model will place some probability mass on occasional greater distances, and hence better accommodate also distant genomes which might otherwise have been considered as different strains.

Some of the resulting probability mass functions 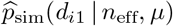 were already shown in Fig 4. In practice, the computations above are done using logarithms and the fact log, to avoid numerical underflow, which can occur whenever *a_i_* ≪ 0. The finite sample size *s* causes some numerical error, but, because the distances are usually small enough that the number of values we need to consider is limited, *s* can be made large enough without too extensive computation, making this error small in general. The above procedure allows computation of the conditional density in Eq 21 for any (*n*_eff_*, μ*), and we can use a Metropolis update for (*n*_eff_, *μ*). We marginalised λ in Eq 21 to improve the mixing of the chain and to be able to use the analytical formula in Eq 22, and in the supplementary material we justify that this algorithm is valid under the assumption that a new λ parameter is sampled only if the corresponding proposed value (*n*_eff_, *μ*) has been accepted.

Whenever a new (*n*_eff_, *μ*)-parameter is proposed, we need to compute *p*_sim_ at this point to check the acceptance condition. This value is also needed when sampling z. However, computing *p*_sim_ on each MCMC iteration as described earlier makes the algorithm slow. Consequently, we instead precompute the values of *p*_sim_ in a dense grid of (*n*_eff_, *μ*)-points which can be done in a parallel manner on a computer cluster. Given the grid values, we use bilinear interpolation to approximate *p*_sim_ at each proposed point 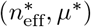. We proceed similarly also with the prior density *p*(*n*_eff_, *μ*|*D*_0_). This approach also allows one to fit the mixture model using different modelling assumptions or different data sets without need to repeat the costly W-F simulations.

Finally, we see that the probability density of *_i_* conditioned on the other variables does not depend on *η_j_, j* ≠ = *i*. Specifically, we obtain 
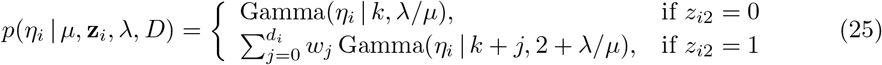
for *i* = 1*,…, N*. Derivation of this result, the formula for the mixture weights *w_j_* and a special algorithm (Algorithm 2) to generate random values from this density are shown in the supplementary material.

The resulting Gibbs sampler is presented as Algorithm 1. It could be alternatively called a Metropolis-within-Gibbs sampler since some of the parameters (*n*_eff_ and *μ*) are sampled using a Metropolis-Hastings step using a proposal density that is denoted as *q*. Because *n*_eff_ is a discrete random variable, (*n*_eff_*, μ*) is a mixed random vector and we cannot use the standard Gaussian proposal. Instead, we consider the distribution 
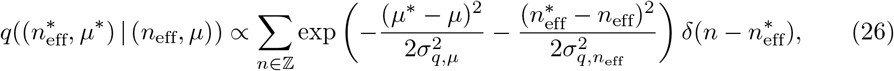
where 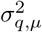 and 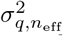 are chosen to produce acceptance probability of the Metropolis step close to 0.25 and *δ*(*·*) is the Dirac delta function. The first element of a random sample from *q* in Eq 26 is an integer, and this proposal is also symmetric. We truncate the tails of *q* with respect to *n*_eff_ to be able to sample the discrete element from *q* efficiently. In practice we then use a proposal *q* that is a mixture density where the components are as in Eq 26 but with different variance parameters 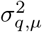 and 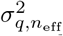 to occasionally propose large steps to increase the exploration of the parameter space.

### Algorithm 1 MH-within-Gibbs sampling algorithm for the mixture model

select an initial parameter *θ*^′^(0) (e.g. by sampling from the prior *p*(*θ*^′^)), proposal *q* and the number of samples *s*

**for** *i* = 1,…, *s* **do**

sample 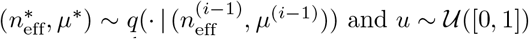 and *u* ∼ *U*([0, 1])

compute *ρ* = min 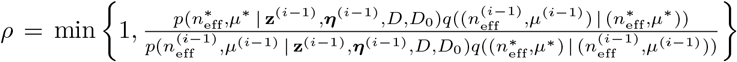 using Eq 21

**if** *ρ* < *u* **then**

set 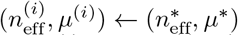

sample λ^*(i)*^ ∼ *p*(η|*μ^(i)^, η^(i−1))^*, λ = λ^(*i*)^ using Eq 22

**else**

set 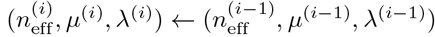

**end if**

**for** *j* = 1,…, *N* do

sample 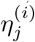 using the Algorithm 2 with *μ* = *μ*^(*i*)^, z = z^(*i*−1)^, = λ = λ^(*i*)^

**end for**

**for** *j* = 1,…,*N* do

sample 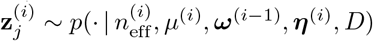 using Eq 18

**end for**

sample *ω*^(*i*)^ ∼ *p*(·|z^(*i*)^, *D*) using Eq 17

**end for**

**return** samples 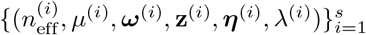

## Posterior predictive distribution

Given a new (future) data point (*d^*^, t^*^*) from a new patient, we would like to compute the probability of whether this case is of the same strain. This can be computed from the posterior of the model fitted to data *D, D*_0_ as follows. We denote the original parameter vector with *θ* as before and additional parameters related to the new data point *D^*^* = {(*d^*^, t^*^*)} as z^*^ ∈ {(1, 0), (0, 1)} and 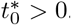. The updated posterior after considering the new data point *D^*^* is then 
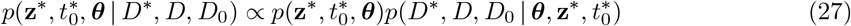

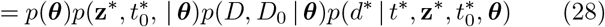

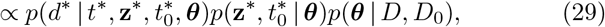
where *p*(*θ*|*D, D*_0_) is the posterior based on our original data *D, D*_0_. We marginalise the set of parameters least contributory to the aim to obtain 
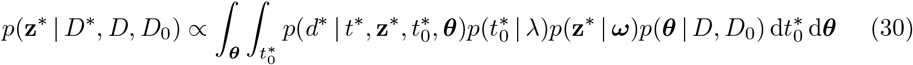

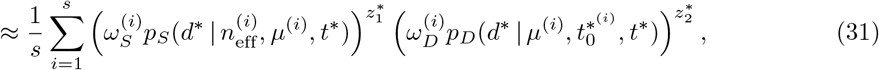
where 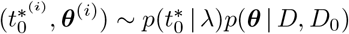 for *i* = 1,…, *s*. The probability of the new measurement point (*d^*^, t^*^*) being of the same strain, based on the previously observed data *D, D*_0_ is obtained from Eq. 31.

## Results

In this section we fit the W-F model to the external data *D*_0_ as discussed in Section ABC inference to update the prior using external data. We then verify that the proposed Gibbs sampling algorithm for fitting the mixture model from Section Bayesian inference for the mixture model is consistent based on experiments with simulated data. Subsequently, we fit the mixture model to the MRSA data and discuss the results. Finally, we assess the quality of the model fit.

### Updating the prior using ABC inference

The ABC posterior based on the external data *D*_0_ and the discrepancy in Eq 8, is shown in Fig 5A. We also repeated the computations so that we omitted a subset, patients A-M, from the analysis i.e. the second summation term in Eq 8 was set to zero. This was done to assess the effect of patients A-M, which have measurements from one time point only, and an unknown time since acquisition. This extra analysis resulted in an ABC posterior approximation shown in Fig 5B. We see that in both cases large parts of the parameter space have been ruled out as having negligible posterior probability. As expected, the posterior distribution based on the subset (Fig 5B) is slightly more dispersed than with the full data *D*_0_ (Fig 5A). Using the full data causes the estimated mutation rate to be slightly greater than with the subset, likely because the model needs to accommodate the higher variability in the patients A-M. In addition, small effective sample sizes (*n*_eff_ < 2000) are less probable based on the full data *D*_0_.

**Fig 5.**
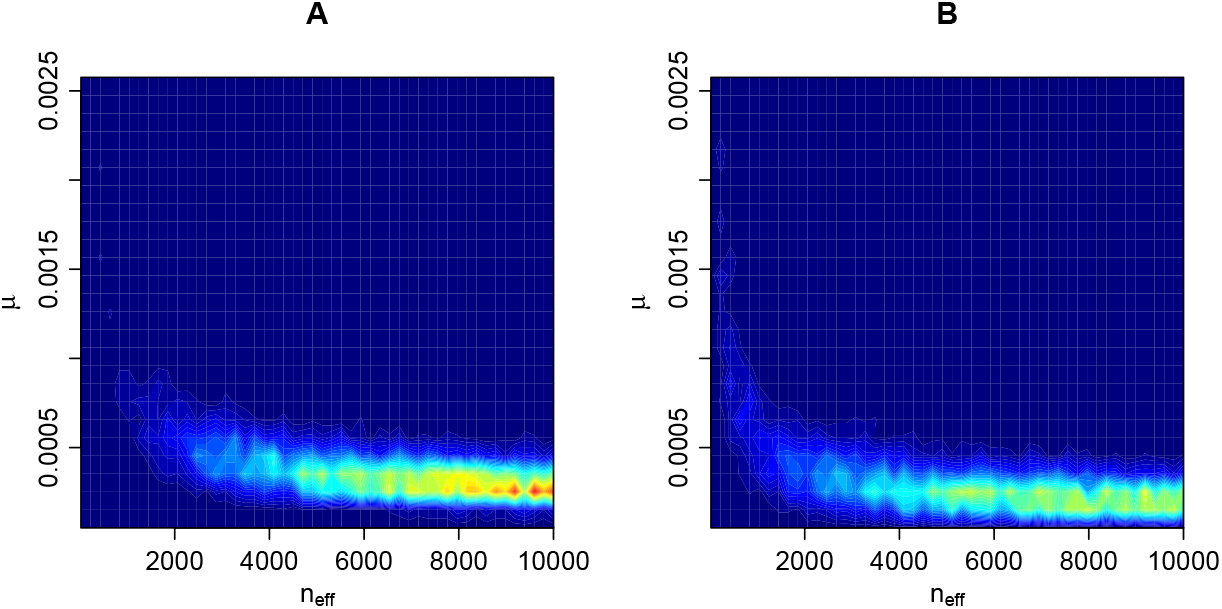
ABC posterior distribution for (*n*_eff_*, μ*). The ABC posterior distribution i.e. the updated prior for parameters (*n*_eff_, *μ*), the effective population size and mutation rate, given data *D*_0_. Panel A shows the result with the full data and panel B the corresponding result with only a subset of the data (see text for details).

Overall, we see that the effective sample size *n*_eff_ cannot be well identified based on the external data *D*_0_ alone. We also see that if the upper bound of the prior density of *n*_eff_ was increased from 10, 000, higher values would likely have non-negligible posterior probability also; however, this constraint will have a negligible impact on the resulting posterior from the mixture model as is seen later. The mutation rate *μ*, on the other hand, is smaller than 0.001 mutations per genome per generation with high probability and cannot be arbitrarily small.

### Validation of the mixture model using simulated data

To empirically investigate the identifiability of the mixture model parameters and the correctness and consistency of our MCMC algorithm under the assumption that the model is specified correctly, we first fit the mixture model to simulated data. We generate artificial data from the mixture model with parameter values similar to the estimates for the observed data *D* from the next section. Specifically, we choose *n*_eff_ = 2, 137, *μ* = 0.0011, *ω_S_* = 0.8, = 0.0001 and we repeat the analysis with various data sizes *N*. We use otherwise similar priors as for the real data in the next section except that, for simplicity, instead of using the prior obtained from the ABC inference, we use a uniform prior in Eq 7. We then fit the mixture model to the simulated data sets to investigate if the true parameters can be recovered (identifiability) and whether the posterior becomes concentrated around their true values when the amount of data increases (consistency).

Results are illustrated in Fig 6. We see that the (marginal) posterior of (*n*_eff_, *μ*) is concentrated around the true parameter value that was used to generate the data (green diamond in the figure). Also, despite the fact that the number of parameters increases as a function of data size *N* (because each data point (*d_i_, t_i_*) has its own class indicator z_*i*_ and time to the most recent common ancestor *t*_0*i*_ parameter), the marginal posterior distribution of (*n*_eff_, *μ*) can be identified and appears to converge to the true value as *N* increases. On the other hand, we cannot learn each *t*_0*i*_ accurately since essentially only the data point to which the parameter corresponds provides information about its value. However, precise estimates of these nuisance parameters are not needed for using the model or obtaining useful estimates of the other unknown parameters as demonstrated in Fig 6.

**Fig 6.**
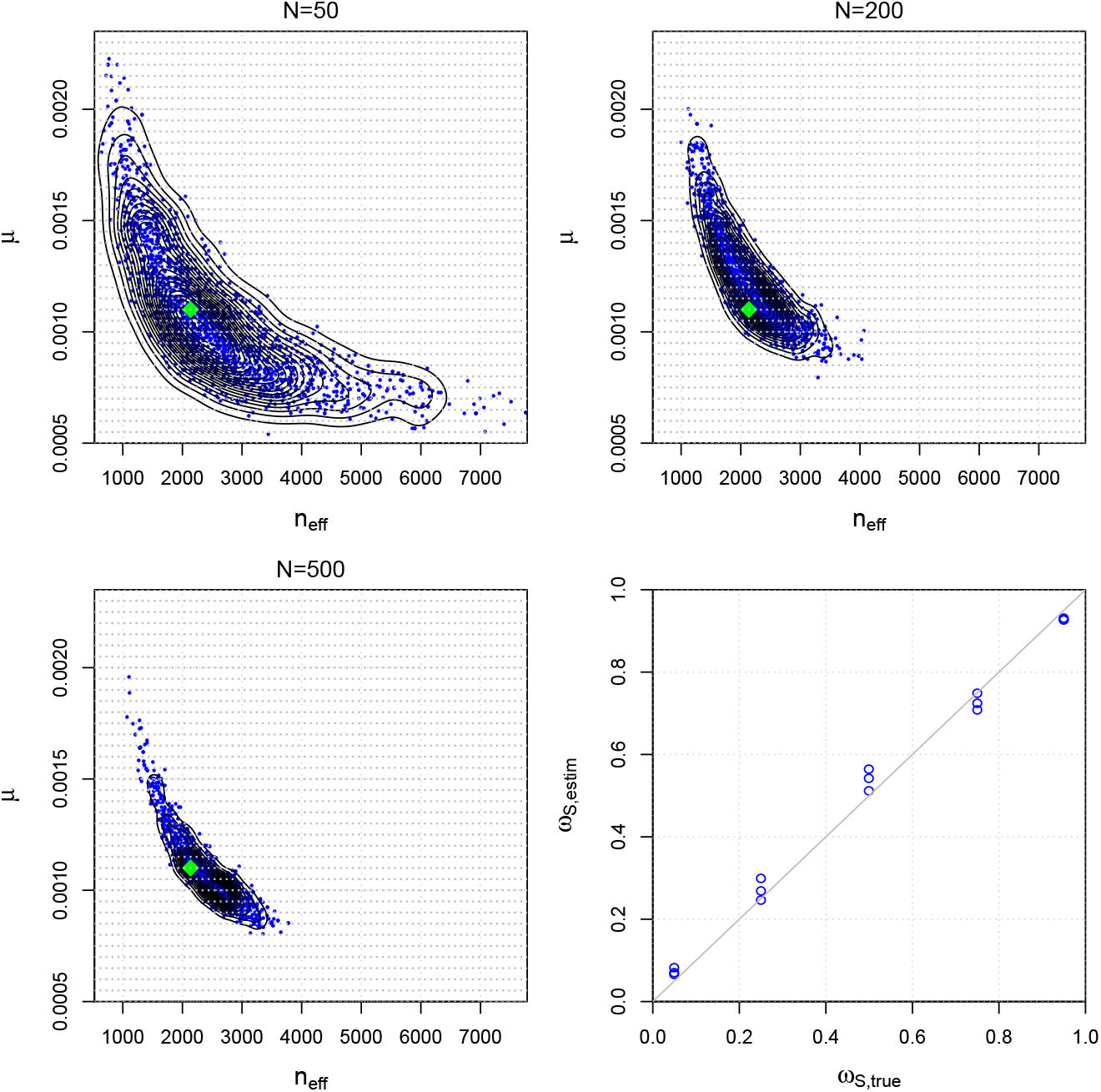
Accuracy and consistency with synthetic data. The first three panels show the estimated posterior distributions for parameters (*n*_eff_, *μ*) of the mixture model using simulated data of different sizes *N*. The green diamond shows the true value used to generate the simulated data and the light grey dots denote the grid point locations needed for numerical computations. The bottom right panel shows the estimated vs. the true *ω_S_* parameter in a set of additional simulation experiments.

The panel in the lower right corner of Fig 6 shows results from an additional simulation experiment where the mixture model is fitted to data generated with different values for the *ω_S_* parameter, which represents the proportion of pairs that are from the same strain. Other than that and the fact that we fixed *N* = 150, the experimental design is the same as above. The results show that the estimated *ω_S_* values generally agree well with the true values. Interestingly, *ω_S_* is slightly overestimated when its true value is close to 0, and slightly underestimated when the true value is close to 1, which may reflect the regularizing effect of the prior, drawing the estimates away from the extreme values. Furthermore, when the true value of *ω_S_* is around 0.5, the variance of the estimate tends to be higher than with *ω_S_* values close to 0 or 1. This observation may be explained by the fact that there are more data points that overlap both mixture model components when *ω_S_* is around 0.5 which makes the inference task more challenging and causes higher posterior variance.

### Analysis of the Project CLEAR MRSA data

The following settings are used to analyse longitudinally-sampled *S. aureus* nares isolates from the control arm of Project CLEAR [23]. We generate 4 MCMC chains, each of length 25, 000, initialized randomly from the prior density, whose first halves are discarded as ∊burn-in”. We use the Gelman and Rubin’s convergence diagnostic in R-package coda and visual checks to assess the convergence of the MCMC algorithm. We use 100 × 100 equidistant grid for numerical computation with the (*n*_eff_, *μ*) values and *s* = 10, 000 in Eq 24. The ABC posterior obtained in Section Updating the prior using ABC inference and visualised in Fig 5A is used as the prior for (*n*_eff_, *μ*).

The parameter vector θ consists of the ’global’ parameters *n*_eff_, *μ, ω, λ*, as well as a large number of nuisance parameters (z and t_0_) related to each data point. The estimated global parameters are presented in Table 1. We also repeated the analysis using a uniform prior on (*n*_eff_, *μ*). While the uniform prior is non-informative about the parameters (*n*_eff_, *μ*), the results are nevertheless surprisingly similar (Table 1). In other words, the additional data *D*_0_ used to update the prior has only a small effect on the estimated parameters of the mixture model. This was unexpected because the data set *D* used to train the mixture model has only one genome per sampled time point, and yet, impressively, the model is able to learn about the parameters (*n*_eff_, *μ*) which effectively define the variability in the whole population. This further demonstrates the robustness of the mixture model to the prior used. We observe, however, that incorporating the prior from the ABC slightly shifts the probability distribution for *n*_eff_ towards larger values, although there is no clear conflict between the two results. For example, as seen in Table 1, the 95% credible interval (CI) for *n*_eff_, [1200, 2200], gets updated to [1300, 2200] when the extra prior information is included.

**Table 1.** Posterior mean and 95% credible interval (CI) for the ’global’ parameters of the mixture model.

| parameter | Informative prior (ABC, data $D_0$) | | Uniform prior | |
| --- | --- | --- | --- | --- |
|  | mean | 95% CI | mean | 95% CI |
| $n_{\text{eff}}$ | 1700 | [1300, 2200] | 1700 | [1200, 2200] |
| $\mu$ | 0.00076 | [0.00060, 0.00092] | 0.00080 | [0.00064, 0.00095] |
| $\omega_S$ | 0.87 | [0.83, 0.91] | 0.88 | [0.83, 0.92] |
| $\omega_D$ | 0.13 | [0.09, 0.17] | 0.12 | [0.08, 0.17] |
| $\lambda (\times 10^5)$ | 7.3 | [5.8, 9.0] | 7.5 | [5.9, 9.3] |

Fig 7 shows the posterior predictive distribution for the probability of the same strain case for a (hypothetical future) observation with distance *d*^*^ and time difference *t^*^*. Blue colour in the figure denotes high probability of the same strain. The corresponding 50% classification curve is (almost) a straight line with a steep positive slope. This is as expected since the same strain model can explain a greater number of mutations when more time has passed. Approximately 20 mutations draws the line between the same strain and different strains cases within the time difference up to 6000 generations. The uncertainty in the classification occurs because there is overlap in the two explanations (around *d*^*^ ≈ 20) and because of the posterior uncertainty in the model parameters θ.

**Fig 7.**
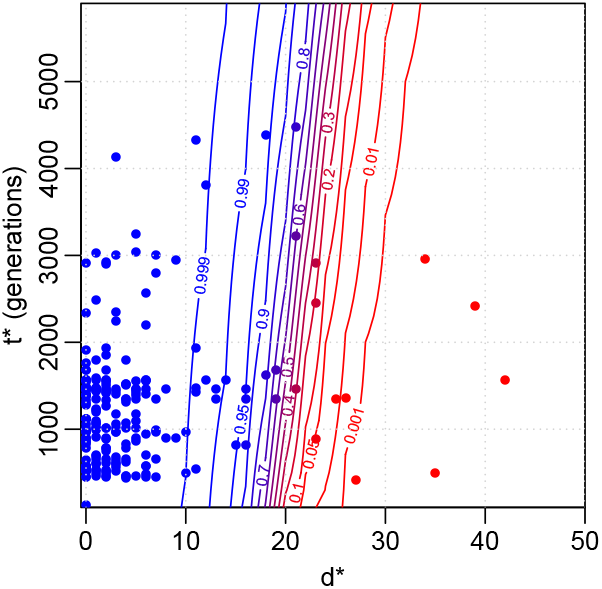
Results for the Project CLEAR MRSA data. Contour plot for same strain probability of a distance *d^*^* and time interval *t^*^* based on the fitted model. The coloured points denote the observations that were used to fit the model. Blue colour indicates large same strain probability. Distances greater than 50 are not shown and are classified as different strains with probability one. 6, 000 generations on the y-axis correspond to approximately one year.

We also analysed explicitly all observed patterns where: 1) two genomes of the same ST from the same patient are interleaved with a missing observation, i.e. the colonization appears to disappear and then re-emerge, and 2) two genomes of the same ST from the same patient are interleaved with an observation of a different ST. The numbers for the two genomes being from the same or different strain in these patterns are shown in Table 2. The credible intervals for the ’same strain’ proportion combine uncertainty from the limited number of samples with the posterior uncertainty of whether a sample is from the same strain or not (see the Supplementary material for further details). From Table 2 we see that approximately 58% of genome pairs in pattern 1) are from the same strain. This is only a little smaller than the same strain proportion when there are no missing observations in between (84%). Therefore, a plausible explanation for most of the missing in-between observations is that in reality the same strain has been colonizing the patient throughout, and the missing observation reflects the limited sensitivity of the sampling, rather than a clearance followed by a novel acquisition. Similarly, even if interleaved with a different ST (pattern 2), the surrounding genomes often, in 63% of cases, appear to be from the same strain. This suggests that in these cases the patient has been colonized by the surrounding strain throughout, and co-colonized by two different STs at the time of observing the divergent ST in the middle.

**Table 2.** The estimated numbers (mean, 95% CI in parenthesis) of cases with genomes in the beginning and in the end of the pattern being from the same or different strain, for three different patterns in the Project CLEAR MRSA data, and the estimated proportion of the same strain cases. same strain/n diff. strain/n same strain prop.

|  | same strain/n | diff. strain/n | same strain prop. |
| --- | --- | --- | --- |
| ST A → ST A | 190(187, 192)/224 | 34(32, 37)/224 | 0.84(0.78, 0.89) |
| ST A $\rightarrow \emptyset \rightarrow \dots \rightarrow$ ST A | 17(16, 19)/29 | 12(10, 13)/29 | 0.58(0.36, 0.76) |
| ST A → ST B $\rightarrow \dots \rightarrow$ ST A | 12(10, 12)/18 | 6(6, 8)/18 | 0.63(0.34, 0.81) |

“ST A → ST A” denotes the case where the ST does not change between two genomes at consecutive samples, “ST A $\rightarrow \emptyset \rightarrow \dots \rightarrow$ ST A” is the pattern 1) where one or more negative samples are seen between the same ST and “ST A → ST B $\rightarrow \dots \rightarrow$ ST A” is the pattern 2) where a sample with different ST is observed between two samples of the same ST. $n$ denotes the number of data points in each alternative.

Finally, we compute acquisition and clearance rates using our model, and compare those to the ones obtained with the common strategy of using a fixed distance threshold. For the purposes of this exposition, we define the acquisition *r*_acq_ and clearance rates *r* clear informally as 
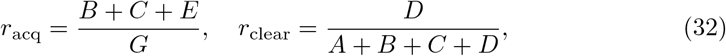
where the quantities *A, B, C, D* and *E* denote the numbers of possible events in consecutive samples (e.g. acquisition, replacement, clearance, or no change) defined in detail in Table 3. Also, *G* is the total number of possible events over the whole data. The quantities *A, B, D* and *E* are random variables that depend on the same/different strain posterior probabilities and, consequently, we also compute the uncertainty estimates for these quantities in Eq 32. Number C is a constant because an observed change of ST always indicates an actual change of ST as well. For cases with one or more negative samples (denoted by ∅) between two positive samples, we do not know when the clearance and acquisition events took place and whether the negative samples are ∊false negatives”. To handle these cases, we parsimoniously assume that a missing observation between two positive samples that are inferred to come from the same strain is a false negative (i.e. that the same strain was present also in the middle, even if it was not detected), and record these events in the groups A-E accordingly. Details on how we unambiguously determine the group for all special cases is provided in the Supplementary material.

The estimated acquisition and clearance rates with 95% credible intervals are shown on the last two lines of Table 3. For comparison, we also computed these rates otherwise similarly but using a fixed distance threshold of 40 mutations, a value used in [10], to determine if two genomes are from the same strain or not. We see that the threshold-based estimates are relatively similar to, and only slightly smaller than the estimates from our model. The explanation for the similarity of summaries such as the acquisition and deletion rates is that, when estimating these quantities across the whole data set, the uncertainty gets averaged out, even if individual data points exhibit a lot of uncertainty regarding whether they are the same strain or not (see Fig 7). Importantly, while being consistent with the previous results, our model bypasses the task of heuristically choosing a single threshold and adds uncertainty estimates around the point estimates, crucial for drawing rigorous conclusions.

**Table 3.** Estimated numbers (posterior means) of different patterns A-E of consecutive samples and the estimated acquisition and clearance rates (mean, 95% CI in parenthesis).

| event | expected number |  |
| --- | --- | --- |
| A: ST A, str X → ST A, str X | 231 | |
| B: ST A, str X → ST A, str Y | 34 | |
| C: ST A → ST B | 45 | |
| D: ST A, str X $\rightarrow \emptyset$ | 104 | |
| E: $\emptyset \rightarrow$ ST A, str X | 21 | |
| rate parameter | post. estimate | threshold-based estimate |
| acquisition rate $r_{\text{acq}}$ | 0.18(0.17, 0.19) | 0.16 |
| clearance rate $r_{\text{clear}}$ | 0.25(0.24, 0.25) | 0.24 |

Above, ST denotes sequence type as before, str denotes the strain and symbol; denotes a negative sample i.e. no bacteria detected.

### Assessing the goodness-of-fit of the model for the Project CLEAR MRSA data

As the last part of our analysis, we use posterior predictive checks to assess the quality of the model, see e.g. [15] for further details. Briefly, this consists of simulating replicated data sets *D*^rep,(*j*)^ from the fitted mixture model and comparing these to the observed data *D* for any systematic deviations. Any discrepancies between the observed and simulated data can be used to criticise the model and understand how the model could be improved. In practice, simulating replicate data is done by simulating a parameter vector *θ^(*j*)^* from the posterior (by using the existing MCMC chain) and simulating a new set of distance-time difference pairs 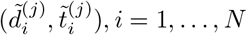 in *D*^rep,(*j*)^ from the model using ^*θ(*j*)*^. To obtain *M* replicates this procedure is repeated for *j* = 1,…, *M*.

Example replicate data sets are shown in Fig 8. Overall, the simulated distances are similar to the corresponding observations. There is a clear peak at *d_i_* = 0, and as the distance is increased the frequency starts to decrease. Occasional large distances (*d_i_* > 20) occur only rarely, in keeping with the observed data. A minor discrepancy is that the fitted model tends to underestimate the frequency of distance zero while small positive distances tend to occur more frequently than observed. This could happen because we estimated the empirical densities *p*_sim_(*d_i_*_1_ *|n*_eff_, *μ*) using a constant time of 6, 000 (i.e. 1 year) since the acquisition (as discussed in Section Bayesian inference for the mixture model), which may lead to a slight overestimation of the distances. To explore the impact of this assumption further, we repeated the analysis so that we computed the densities *p*_sim_(*d_i_*_1_ *| n*_eff_, *μ*) at a constant time of 1, 000 generations. However, the mismatch did not disappear completely and the estimated mutation rate increased as a result to compensate for the occurrence of greater distances, in disagreement with the prior density from the ABC analysis and data *D*_0_. We thus believe that the current model is adequate.

**Fig 8.**
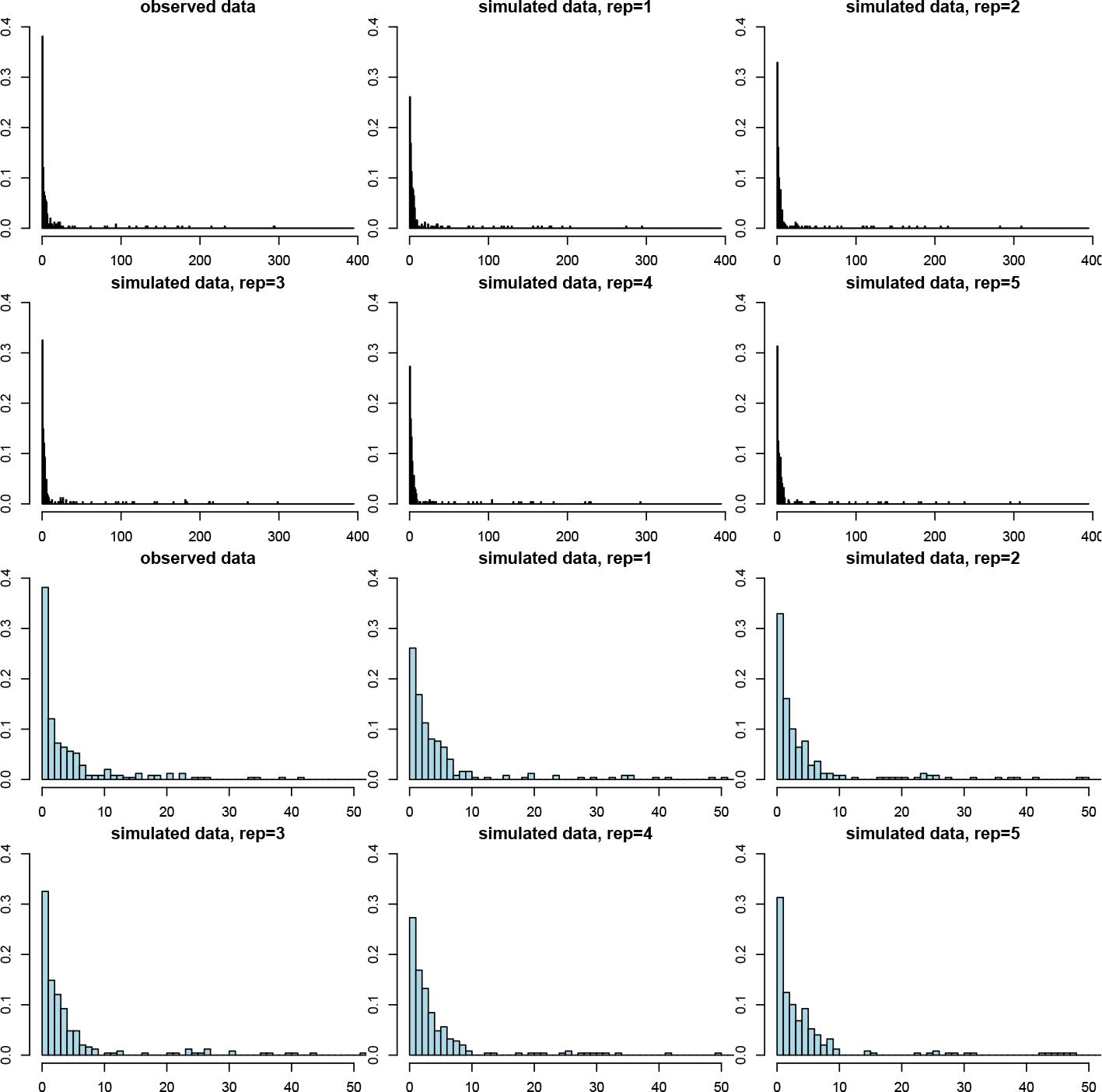
Model validation using posterior predictive checking. The histogram in the upper left corner shows the observed distance distribution in the Project CLEAR MRSA data, the other figures in the top two rows show the corresponding distances in replicate data sets simulated from the fitted model. The bottom two rows show the same histograms zoomed to range [0, 50]. The replicate data sets look overall similar to the observed data, demonstrating the adequacy of the model. However, the amount of zero distances is underestimated and the frequencies of small positive distances tend to be slightly overestimated.

## Discussion

We presented a new model for the analysis of clearance and acquisition of bacterial colonization, which, unlike previous approaches, does not rely on a heuristic fixed distance threshold to determine whether genomes observed at different times points are from the same or different acquisition. Fully probabilistic, the model automatically provides uncertainty estimates for all relevant quantities. Furthermore, it takes into account the variation in the time intervals between pairs of consecutive samples. Another benefit is that the model can easily incorporate additional external data to inform about the values of the parameters. To fit the model, we developed an innovative combination of ABC and MCMC, based on an underlying mixture model where one of the component distributions was formulated empirically by simulation.

We demonstrated the model using data on *S. aureus* genomes sampled longitudinally from multiple patients. Our analysis provided evidence for occasional co-colonization and identified likely false negative samples. The output of the model consists of the same vs. different strain probability for any pair of genomes, and, by using this information to decide (probabilistically) when and where the colonizing strain had changed, the acquisition and clearance rates were easy to calculate. Estimates of these parameters were found to be in agreement with previous estimates derived using a fixed threshold, but now we were able to provide confidence intervals, essential for drawing rigorously supported conclusions. We believe such analyses are common enough that our method should be useful for many, and, consequently, we provide it as an easy-to-use R-code. The code includes tools for both the ABC-inference to incorporate external data of distance distributions between multiple samples at a given time point (or two time points), and the MCMC-algorithm. We note that our method does not assume recombination, which was not relevant with the present data. If this is an issue, we recommend removing recombinations by preprocessing the genomes with one of the standard methods [30–32]. While our analysis demonstrated that the external data may reduce uncertainty in the resulting posterior, we also saw that the method may work without such data. In the latter case the input is simply a list of distance-time difference pairs for genomes sampled from the same patient at consecutive time points, and it is sufficient to run the MCMC, which is efficient and fast in typical cases.

A central component of our approach is a model for within-host variation, required to determine how much variation can be expected if the genomes at different time points have evolved from the same strain obtained in a single acquisition. We selected for this purpose the basic Wright-Fisher model assuming constant population size and mutation rate with the understanding that these assumptions are expected to be violated to some extent in any realistic data set, but the benefits of simplicity include robustness of the conclusions to prior distributions and identifiability of the parameters from the available data. More complex models have been fitted to the distance distributions (our external data *D*_0_), assuming the population size first increases and then decreases [13]. However, our model can fit the same data with fewer parameters, which justifies the simpler alternative. Furthermore, the constant population size may also be seen as a sensible model for persistent colonization. An interesting future research question is what additional data should be collected in order to be able to fit one of the possible extensions of the basic model. Another direction that we are currently pursuing is to extend the model to cover genomes sampled from multiple body sites.

## Supporting information

**S1 File. Derivations and further details of the model**. We provide some further derivations and details related to our MCMC algorithm. To guide the selection of prior hyperparameters, we also derive the explicit prior distribution and some of its summaries for the parameter t_0_ and the mean and variance for the prior predictive distribution for the distance. We also describe further details on computing the acquisition and clearance rates.

## Acknowledgments

We acknowledge the computational resources provided by Aalto Science-IT project.

